# invertiaDB: A Database of Inverted Repeats Across Organismal Genomes

**DOI:** 10.1101/2024.11.11.622808

**Authors:** Kimonas Provatas, Nikol Chantzi, Michail Patsakis, Akshatha Nayak, Ioannis Mouratidis, Georgios A. Pavlopoulos, Ilias Georgakopoulos-Soares

**Affiliations:** Institute for Personalized Medicine, Department of Biochemistry and Molecular Biology, The Pennsylvania State University College of Medicine, Hershey, PA, USA; Huck Institute of the Life Sciences, Pennsylvania State University, University Park, PA, USA; Institute for Fundamental Biomedical Research, BSRC “Alexander Fleming”, Vari 16672, Greece

## Abstract

Inverted repeats are repetitive elements that can form hairpin and cruciform structures. They are linked to genomic instability, however they also have various biological functions. Their distribution differs markedly across taxonomic groups in the tree of life, and they exhibit high polymorphism due to their inherent genomic instability. Advances in sequencing technologies and declined costs have enabled the generation of an ever-growing number of complete genomes for organisms across taxonomic groups in the tree of life. However, a comprehensive database encompassing inverted repeats across diverse organismal genomes has been lacking. We present InvertiaDB, the first comprehensive database of inverted repeats spanning multiple taxa, featuring repeats identified in the genomes of 118,070 organisms across all major taxonomic groups. The database currently hosts 30,067,666 inverted repeat sequences, serving as a centralized, user-friendly repository to perform searches, interactive visualization, and download existing inverted repeat data for independent analysis. invertiaDB is implemented as a web portal for browsing, analyzing and downloading inverted repeat data. invertiaDB is publicly available at https://invertiadb.netlify.app/homepage.html.

## Introduction

The right-handed double helix DNA structure, also known as the canonical B-DNA structure, was originally described by Watson, Crick, Wilkins and Franklin. Several alternative DNA conformations have since been identified (Ghosh and Bansal 2003; Kaushik et al. 2016; Choi and Majima 2011). These non-canonical DNA structures that do not conform to the standard B-DNA form are termed non-B DNA. Such conformations include hairpin and cruciform structures, which can form at inverted repeat (IR) sequences (Bikard et al. 2010; Buisson et al. 2019; Lu et al. 2015; Bacolla et al. 2016). IRs are composed of two sequence parts, one of which is the reverse complement of the other, separated by an intervening spacer sequence. IRs undergo intrastrand base pairing to form hairpin or cruciform structures, in which the two complementary arms hybridize and the spacer loop remains single-stranded.

The formation of hairpin and cruciform structures can be facilitated by negative supercoiling, often associated with DNA transcription and replication (Azeroglu et al. 2014; Rosche, Trinh, and Sinden 1995). The biophysical properties of IRs, including the spacer and arm lengths and their GC content influence the likelihood of structure formation. Increased GC content in the IR arms is linked to increased hairpin stability (Buisson et al. 2019; Woodside et al. 2006; SantaLucia 1998), whereas mismatches in the arms reduce the likelihood of structure formation (Rentzeperis et al. 2002; Nag and Petes 1991; Nasar, Jankowski, and Nag 2000; Lobachev et al. 1998; Sinden et al. 1991). Additionally, longer arms are linked to increased hairpin formation likelihood and stability (Woodside et al. 2006), while shorter spacer lengths are favorable to hairpin and cruciform structure formation (Woodside et al. 2006). Specifically, cruciforms favor shorter spacer lengths, whereas hairpins favor short spacers but with spacer lengths above four base pairs due to steric constraints (Woodside et al. 2006; Bi and Liu 1996).

IRs are over-represented in organismal genomes relative to the expected random distribution and show an inhomogeneous genomic distribution with enrichment hotspots (Strawbridge et al. 2010). They are particularly enriched in promoters and near transcription termination sites and can regulate gene expression (Georgakopoulos-Soares, Victorino, et al. 2022; Brázda et al. 2020). In prokaryotes, IRs can drive rho-independent transcription termination (Kingsford, Ayanbule, and Salzberg 2007; von Hippel 1998). At replication origins, IRs are involved in replication initiation (Pearson et al. 1996), and in certain transposons, they are found flanking the internal sequence (Fattash et al. 2013). Additionally, several proteins can bind preferentially to hairpin and cruciform structures (Bowater, Bohálová, and Brázda 2022).

Hairpins and cruciforms are associated with increased genomic instability in both prokaryotes and eukaryotes (Gordenin et al. 1993; Lobachev, Rattray, and Narayanan 2007; Nag and Kurst 1997; Lobachev et al. 1998; Butler, Gillespie, and Steele 2002; Achaz et al. 2003; Leach 1994; Tanaka et al. 2002; Zhou, Akgūn, and Jasin 2001; Lindsey and Leach 1989; Georgakopoulos-Soares et al. 2018; Georgakopoulos-Soares, Victorino, et al. 2022; Lu et al. 2015). The human genome is depleted of perfect IRs with long arms, which are highly unstable (Bastos et al. 2023; Wang and Leung 2006; Buisson et al. 2019). IRs are recombination and rearrangement hotspots across organisms (Achaz et al. 2003; Butler, Gillespie, and Steele 2002) and artificial introduction of IRs with long arms in eukaryotic cells leads to thousands of times higher recombination and deletion rates than expected (Lobachev et al. 1998). In human cancers, IRs are highly enriched in somatic mutations across mutation categories and this excess can confound statistical models aimed at identifying driver mutations (Zou et al. 2017; Nik-Zainal et al. 2016; Georgakopoulos-Soares et al. 2018). Specifically, off-target APOBEC mutagenesis is often directed at hairpin structures, causing an excess of mutagenesis at inverted repeats, particularly at the single-stranded loop (Buisson et al. 2019).

Multiple bioinformatic tools have been developed to systematically identify IRs (Brázda et al. 2016; Rice, Longden, and Bleasby 2000; Ye et al. 2014). Additionally, a previous database named Non-B DB encompasses IRs found in five organisms (Cer et al. 2013). However, to date, no database is available to catalog IRs across thousands of organismal genomes, perform dynamic searches, visualize, and download IRs in different genomes.

Here we present invertiaDB, the first IR database across organismal genomes. The database currently hosts 30,067,666 IR sequences found across 118,070 organismal genomes, and serves as a centralized user-friendly web portal to perform searches, interactive visualizations, and download existing IR data for further analysis. This resource will be valuable for researchers across various disciplines, studying the functional roles and genomic instability of inverted repeats.

## Materials and Methods

### Data collection

The organismal genomes present in the database were downloaded from the GenBank and RefSeq databases on 2024-03-21 (O’Leary et al. 2016; Benson et al. 2013) and included all complete organismal genomes. Duplicate assembly accessions were filtered out. Gene annotation files in the form of GFF files were downloaded for each genome from the same source using genome_updater from https://github.com/pirovc/genome_updater. Coordinates for genes, exons, and CDS regions were derived using BEDTools (Quinlan and Hall 2010) and with custom awk, bash, and Python scripts. These coordinates were then examined for IR densities across organismal genomes. The organismal genome database consists of 49,197 Bacteria, 67,725 Viruses, 456 Eukaryota and 687 Archaea.

### Identification of IRs in organismal genomes

IRs with arm lengths equal to or longer than ten bps, spacer lengths less than nine bps, and without mismatches in the arms were used as described in (Chantzi et al. 2024). For IR detection, a modified version of the non-B gfa package was developed and wrapped in a Python program (Cer et al. 2013). The subprocess call to the C script utilized all necessary skip flags to extract only the IR sequences present in each organismal genome. A custom script was used to manually transform the data into a processable tabular format and extract the corresponding arm and spacer sequences along with their corresponding lengths based on the coordinates of the IR sequence. Additionally, the extracted IR sequences were filtered post-extraction, to contain IR sequences with at least 10bp arm length and at most 8bp spacer length. The detected IRs were then systematically validated with custom scripts.

### Database design and data pipelines

The backend of the application is built using Flask, a Python-based web framework, and is served through a reverse proxy to manage internal port access. The data layer leverages DuckDB in read-only mode, optimizing for fast, highly compressed, and secure Online Analytical Processing (OLAP). The DuckDB database driver operates in-process within the Flask app’s memory, with all data consolidated into a single DuckDB file. Currently, the stored inverted repeats occupy 3GB, organized as a vectorized columnar database, accessible through SQL syntax. As of database design, metadata were extracted from the NCBI database for each of the 118,070 assembly genomes that were analyzed for inverted repeats and enriched with the inverted repeats filename to facilitate joint operations between the two. Then, using joins between the metadata and the inverted repeat genomic data, the metadata table was enriched with the inverted repeat statistics (**Supplementary Figure 1)**. We used a custom pipeline to scan the NCBI database for GFF annotations of the 118,070 genomes and we imported them into the database to implement complex queries such as most dense genes in overlapping inverted repeats. Finally, we used various scripts to curate the data and to convert them to multiple formats (Parquet, CSV, JSON, BED), while grouping them into the three domains of life and viruses and providing them for user download.

### Web-interface

The front-end of invertiaDB is built using HTML, CSS, and JavaScript. The application is deployed on a standard Google Cloud Compute Engine, featuring a JavaScript-based front-end app that is served via NGINX operating in web server mode (**Supplementary Figure 2**). A comprehensive full-stack web application was developed to enable user-friendly access, in-depth analysis features, and visualization of the integrated data. To provide in depth understanding of the IR data, dynamic and interactive graphs were also created using custom HTML and Chart.js. Additionally, pre-built components from the Bootstrap 5 and Bootstrap 5 Datatables libraries were utilized.

### Database overview and functionality

#### Database contents and usage

InvertiaDB features comprehensive integration and curation of IRs in 118,070 assemblies, of organisms in the domains of life and viruses, in a user-friendly, interactive web interface (**Figure 1a-b**). Out of all the examined viral genomes, 33,142 did not contain any IRs, while only 1 bacterial genome did not contain any IR sequences. Interestingly, viral genomes displayed greater heterogeneity in their respective IR populations, while bacterial genomes were found to have an abundance of IR sequences. IR sequences can be filtered based on their biophysical properties including their spacer and arm lengths, sequence motifs that the user searches for, and IR metadata such as density, the total IR length, their nucleotide composition and genic overlaps (**Figure 1a-b**). The database includes multiple features to explore, search and download IRs from each organism in Parquet, CSV, JSON and BED formats.

**Figure 1:**
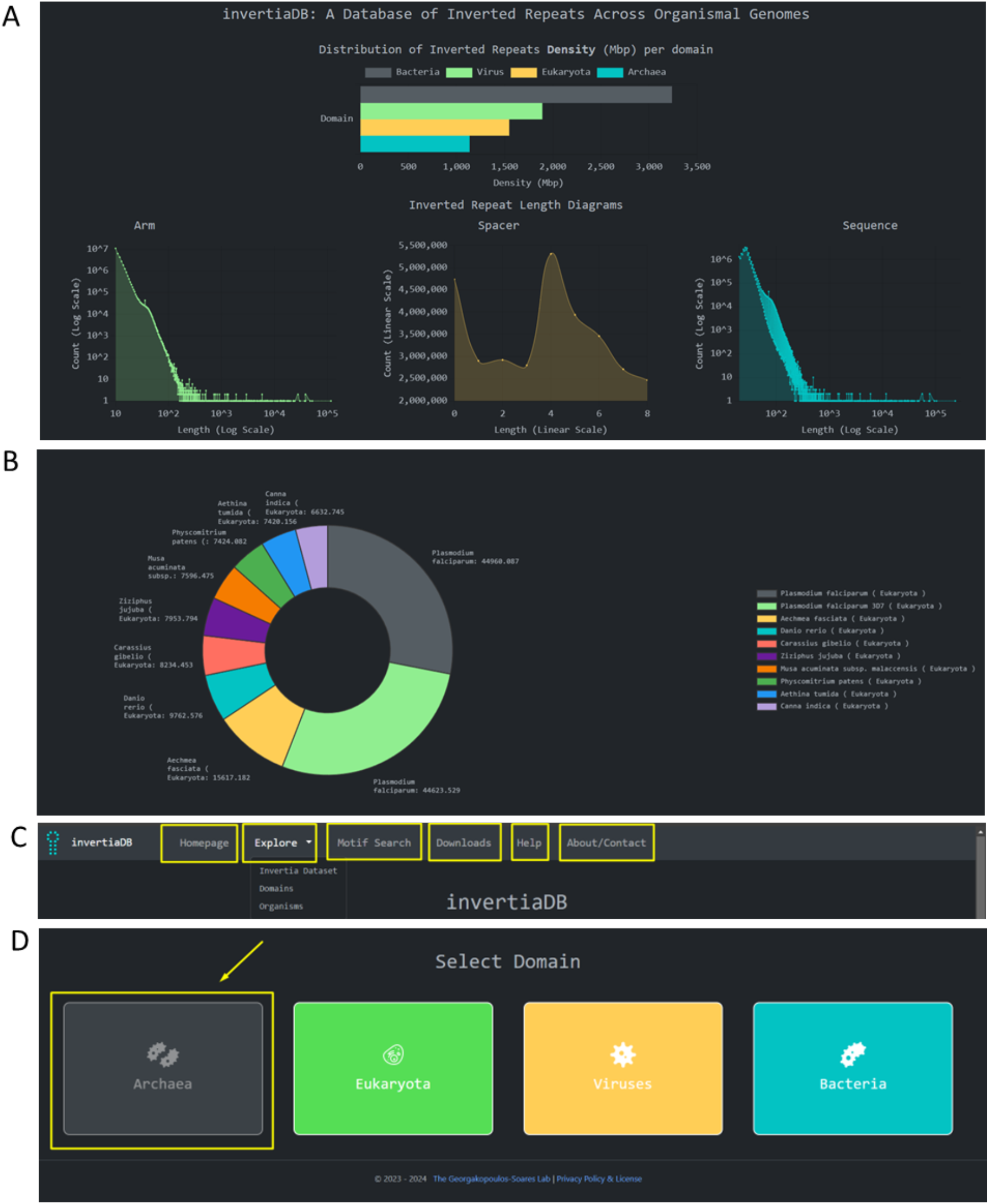
InvertiaDB Homepage, Navigation Bar and Domain selection. **A**. invertiaDB Homepage showing domains of life bar chart in decreasing order of inverted repeat density, and arm, spacer and sequence lengths distribution in line charts. **B**. Doughnut chart of top ten most dense organisms in inverted repeats for each domain of life. **C**. Navigation bar of invertiaDB showcasing each possible option. **D**. Domain page that leads to the aggregated analysis of inverted repeats per domain of life and Viruses.

The top navigation bar is composed of five interactive pages, namely the Homepage, the Explore page, the Downloads page, the Help page and the About/Contact page tabs, which enable the navigation across the different parts of the database (**Figure 1c**). The Explore page is further split into the Invertia Dataset, the Domains page and the Organisms page (**Figure 1c**), with each of these options navigating the user to subsequent pages for IR explorations. The Domain page provides aggregations of IRs by the domain of life, in Archaea, Eukaryota, Bacteria as well as in Viruses (**Figure 1d**).

#### Invertia Dataset page

Upon accessing the Explore → Invertia Dataset page, users are presented with a table of assemblies featuring advanced querying capabilities for exploring IRs across various genomes (**Figure 2a**). Quick searches can be performed on NCBI Taxonomy IDs, GenBank/Reference genome accessions, or species names. Complex queries can be performed using combinations of the available metadata columns provided by the Advanced Filters feature (**Figure 2b**), and users may project all columns from the NCBI database metadata. Multiple genomes can be selected for downloading IR annotations in Parquet, CSV, JSON, or BED formats (**Figure 2b**). Upon clicking the assembly button that is shown as the first column the user navigates to an analysis of the assembly displaying the inverted repeat and genome metadata (**Figure 2c**). The metadata includes links to the established publicly available databases of the ENA Browser (Leinonen et al. 2011) and the NCBI Genome Browser (Pruitt et al. 2009). We generated a visualization depicting the occurrences of inverted repeats in the organism, focusing on the arm-to-spacer ratio. This allows for an assessment of whether the structure is arm- or spacer-dominant, as some organisms exhibit longer spacer/loop lengths relative to arms/stems.

**Figure 2:**
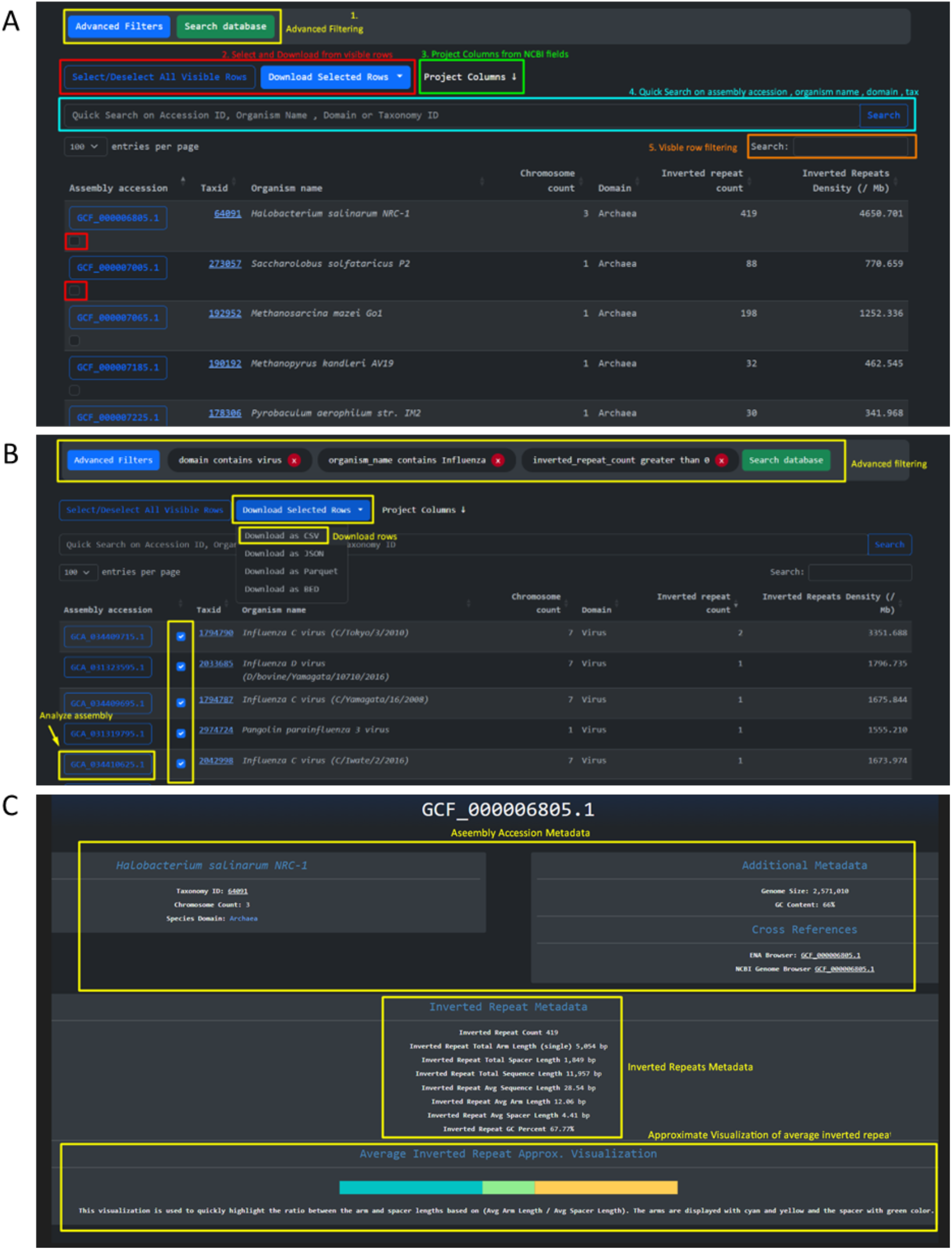
Invertia Dataset Exploration page along with example usage and assembly analysis pages. **A**. Invertia Dataset explore page showcasing 1. Advanced Filtering, 2. Downloading, 3. Projecting NCBI Columns, 4. Quick Search and 5. Row filtering features **B**. Example usage for Influenza Virus containing inverted repeat structures **C**. Assembly accession analysis page, containing metadata and visualizations of genome and inverted repeat patterns.

#### Analysis and visualization pages

The invertiaDB website features interactive bar plots, donut charts, tables, and drop-down menus, allowing users to select and analyze IRs across various assemblies. On the assembly analysis page on inverted repeats, the user also encounters a line chart displaying the length distribution of IRs per arm, spacer, and sequence (**Figure 3a**), along with three donut graphs that showcase respective nucleotide compositions for the three sequences, offering insight into the structural characteristics (**Figure 3a**). A table listing the IRs can be found within the organism’s assembly, with the ability to apply fuzzy search across various columns, enabling filtering of IR data (**Figure 3b**). Finally there are gene-specific analyses with two functionalities: (1) the ability to search for IRs that overlap specific genes by gene locus tag, and (2) a table identifying genes with the highest IR density, calculated by dividing the total size of IRs overlapping the gene divided by the gene’s size in megabase pairs (Mbp) (**Figure 3c**). By default the top ten genes with highest intragenic IR density are displayed in IR bp per megabase.

**Figure 3:**
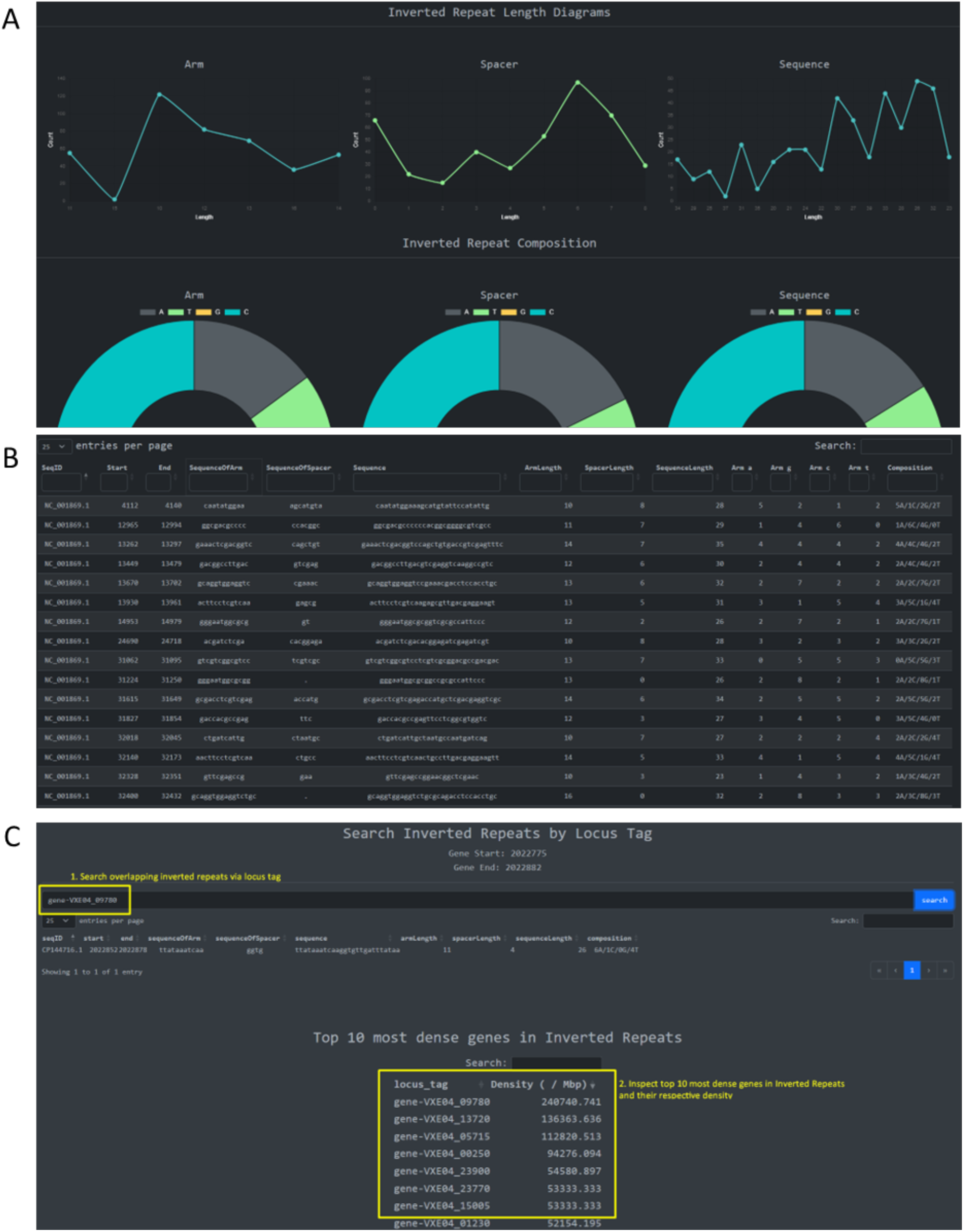
Assembly accession analysis page showing inverted repeat graphs, raw data tables and gene functionalities. **A**. Line chart of length distribution of arm, spacer and sequence along with nucleotide composition of arm, spacer and sequence. **B**. Table of inverted repeats contained in the organism’s assembly with the ability to fuzzy search each column. **C**. 1. Search inverted repeats overlapping specific gene by gene locus tag. 2. Table of most dense genes in inverted repeats by formula: total inverted repeat base-pairs overlapping a gene over gene size in Mbp.

#### Domains and Organisms pages

By accessing the Explore → Domains page the user is redirected to the aggregated IR data in the three domains and viruses. When the user selects a specific domain or Viruses, they are presented with the statistics of average IR count and density across the assemblies, the average spacer, arm and sequence length across assemblies and for individual assemblies in a searchable table (**Figure 4a**). Upon accessing the Explore → Organisms page the results of the Invertia Dataset are arranged by organism name instead of assembly ID (**Figure 4b**). The user can analyze and compare the IR data for multiple assemblies of the same organism when multiple assemblies are present (**Figure 4b-c**). Additionally, by navigating to the individual organism’s page, aggregated IR statistics and metadata are provided, and the user has the option to download these in different file formats (Parquet, CSV, JSON, or BED formats) (**Figure 4c**).

**Figure 4:**
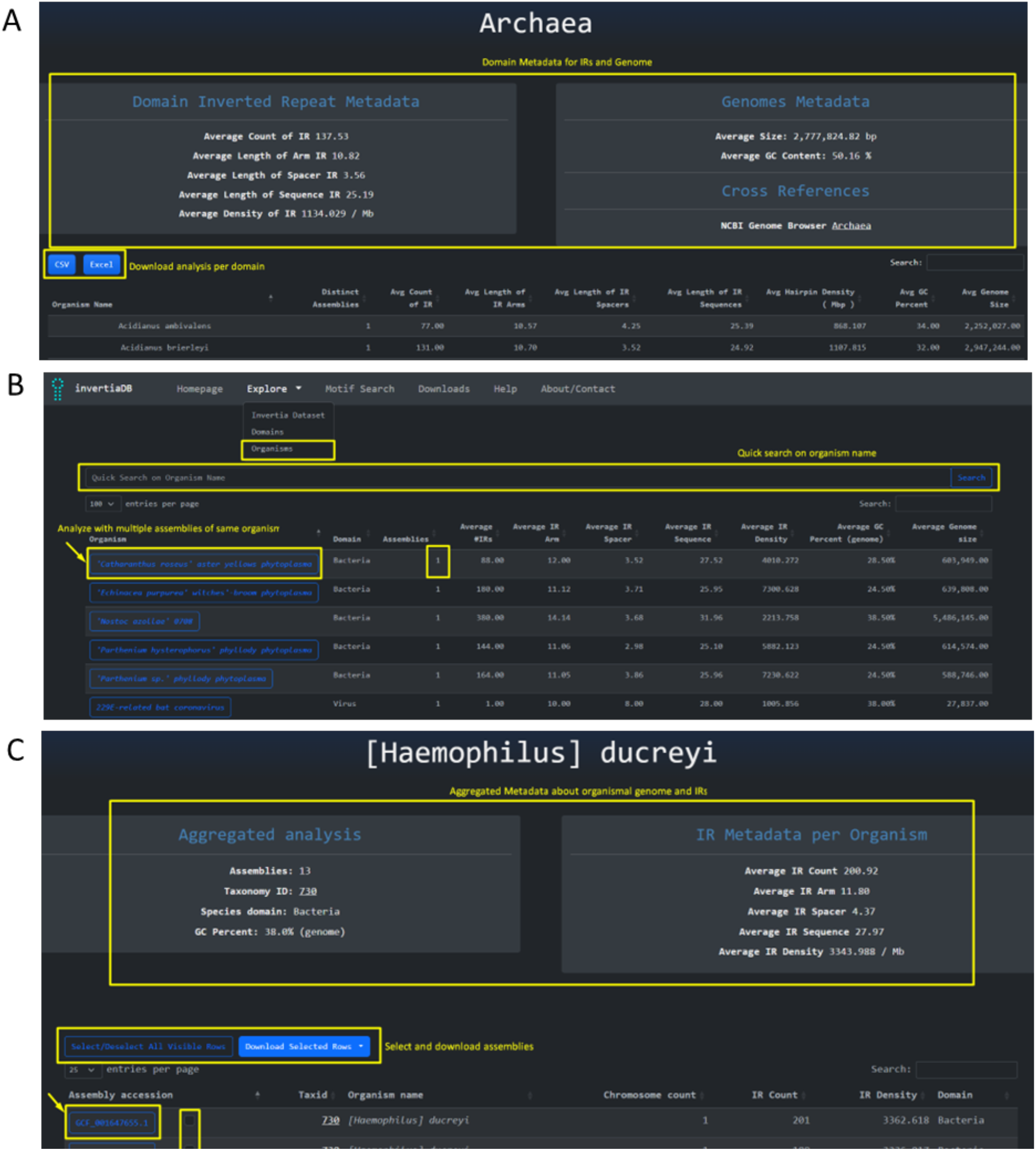
Domain selection and domain analysis along with organism search page and aggregated analysis page. **A**. Analysis of the Archaea domain where each assembly is aggregated per organism along with metadata and the ability to download **B**. Search unique organisms to perform aggregated analysis on their assemblies. **C**. Aggregated analysis (metadata, inverted repeat metadata, file download, file inspection) for specific organisms.

#### Motif search options

The Motif search page enables the search of motifs within IRs or their sub-compartments. There are multiple options to customize the search; these include: 1) String sequence search using standard string filters. The standard string operations that can be used are: equals, contains, starts with, ends with and they can apply sequence wise or granularly on specific regions such as left/right arm, spacer, and both arms. 2) String length search implementing different comparators including “equals”, “greater than”, “less than”, “greater or equal” and “less or equal”. These filters can also apply on different IR regions (**Figure 5a**). After the search filters are applied a metadata search is performed to fetch the different files and the unique organisms that the returned results belong to (**Figure 5b**). 3) Users can also search on the forward, reverse and both strands of the sequences as well for advanced filtering 4) There is also the added option to limit the number of results displayed (**Figure 5c**).

**Figure 5:**
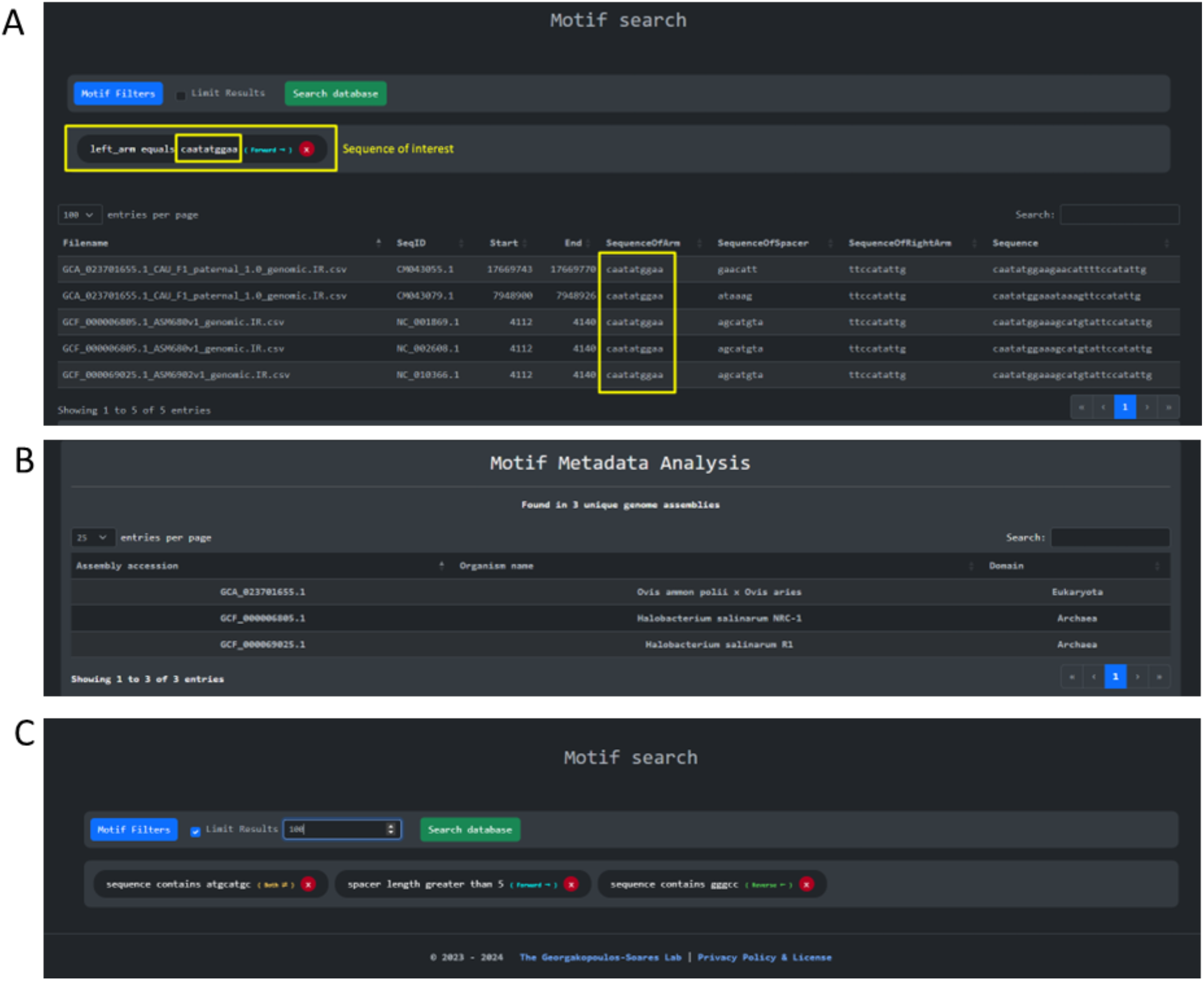
Motif search capabilities. **A**. Example of motif search in the left arm using a single equality filter **B**. Metadata about the motif hits, showing the files and the unique organisms where the motif was found. **C**. Using multiple filters and searching on multiple strands showcasing a more advanced use case.

## Discussion

Here, we have developed invertiaDB, the first centralized and comprehensive IR database spanning all major taxa across the tree of life. The website is user-friendly, interactive and provides various features, including search and filtering tools, dynamic tables that allow for querying and sorting, visualizations and data downloads for in-depth, independent analysis. As the number of available organismal genomes continues to grow, we plan to incorporate them in our database with regular updates.

IRs are a highly dynamic DNA element that can form hairpins and cruciform structures. They are associated with a plethora of functions, but are also linked to increased genomic instability (Gordenin et al. 1993; Lobachev, Rattray, and Narayanan 2007; Nag and Kurst 1997; Lobachev et al. 1998; Butler, Gillespie, and Steele 2002; Achaz et al. 2003; Leach 1994; Tanaka et al. 2002; Zhou, Akgūn, and Jasin 2001; Lindsey and Leach 1989; Georgakopoulos-Soares et al. 2018; Georgakopoulos-Soares, Victorino, et al. 2022). The invertiaDB database enables the systematic examination of the functional roles of IRs, including those in gene regulation, replication, transposition, genome organization and gene and genome evolution among others (Brázda et al. 2020; Georgakopoulos-Soares, Chan, et al. 2022; Georgakopoulos-Soares, Parada, and Hemberg 2022). The integration of biophysical properties, including the spacer and arm lengths, and the nucleotide composition of each IR allows for the study of these parameters and the impact of IR stability to its functional roles and genomic instability. Finally, our database could be utilized by researchers who are interested in examining DNA repair systems associated with hairpin and cruciform structures, which often differ between organisms belonging to different taxonomies (Vasquez and Wang 2013; Pitcher, Brissett, and Doherty 2007).

We envision that invertiaDB will be adopted by researchers around the world to explore the roles and applications of IRs. invertiaDB will enable the systematic exploration of IRs and advance our understanding of their broader roles and impact on organismal evolution and biological functions.

## Code and Data Availability

The invertiaDB dataset can be found in Zenodo with a stable version https://zenodo.org/records/13856709.

## Author Contributions

K.P., N.C., and I.G.S conceived the study. N.C., and K.P. wrote the code, generated the visualizations and performed the analyses with help from M.P., A.N., I.M., G.P, and I.G.S. K.P. developed the database. I.G.S. supervised the project and provided resources. K.P., and I.G.S., wrote the manuscript with help from all authors.

## Declaration of interests

The authors declare no competing interests.

## Funding

This work is supported by the National Institute of General Medical Sciences of the National Institutes of Health under award number R35GM155468. G.A.P. was funded by Fondation Sante; Onassis Foundation; Hellenic Foundation for Research and Innovation (H.F.R.I) under the call ‘Greece 2.0 - Basic Research Financing Action, sub-action II, Grant ID: 16718-PRPFOR; Program ‘Greece 2.0, National Recovery and Resilience Plan’, Grant ID: TAEDR-0539180.

## Declaration of interests

The authors declare no competing interests.

## Supplementary Material

**Supplementary Figure 1:**
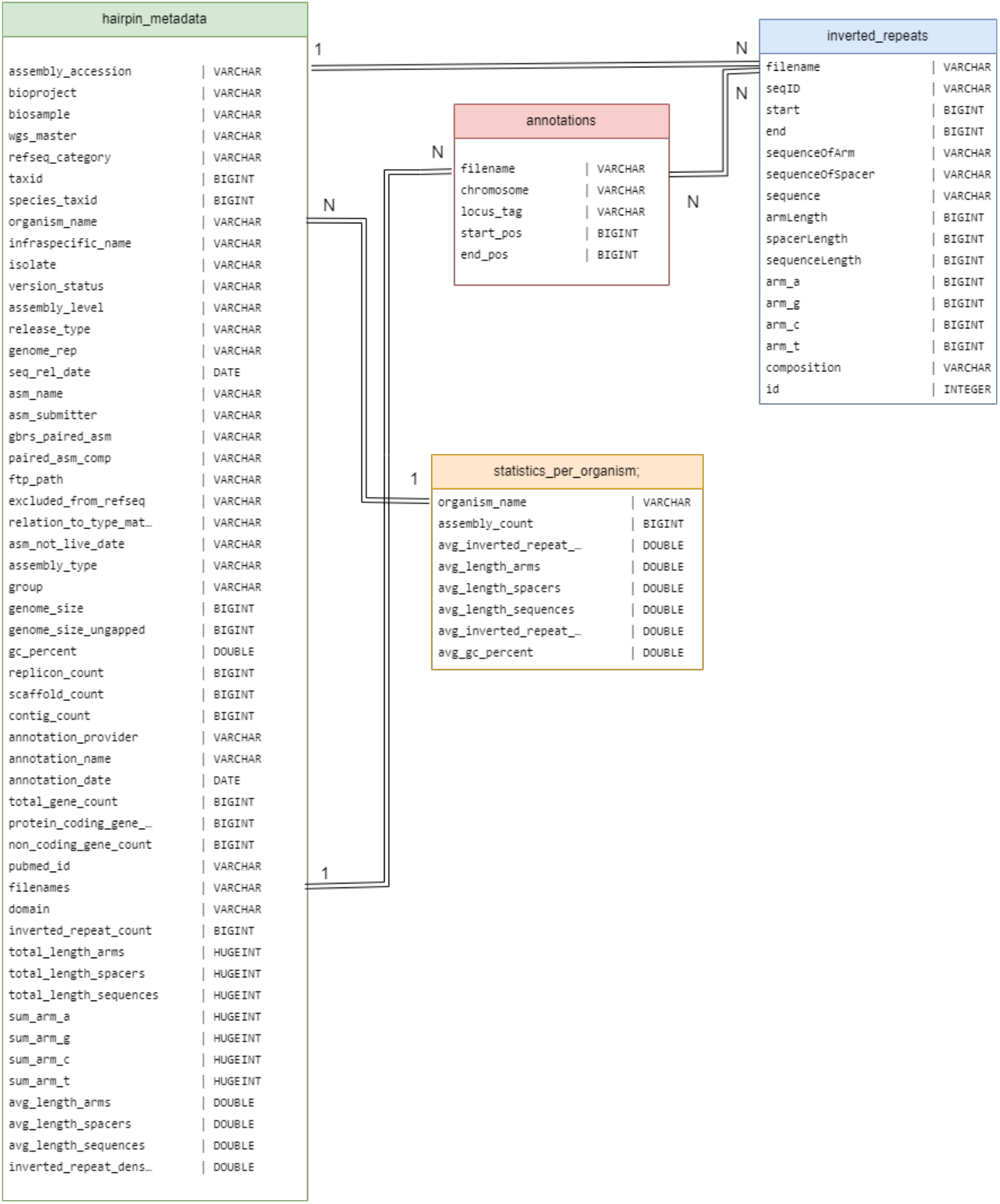
Database Entity Relationship Diagram. This diagram is used to describe the column content of each database table and the cardinality relationships between entities.

**Supplementary Figure 2:**
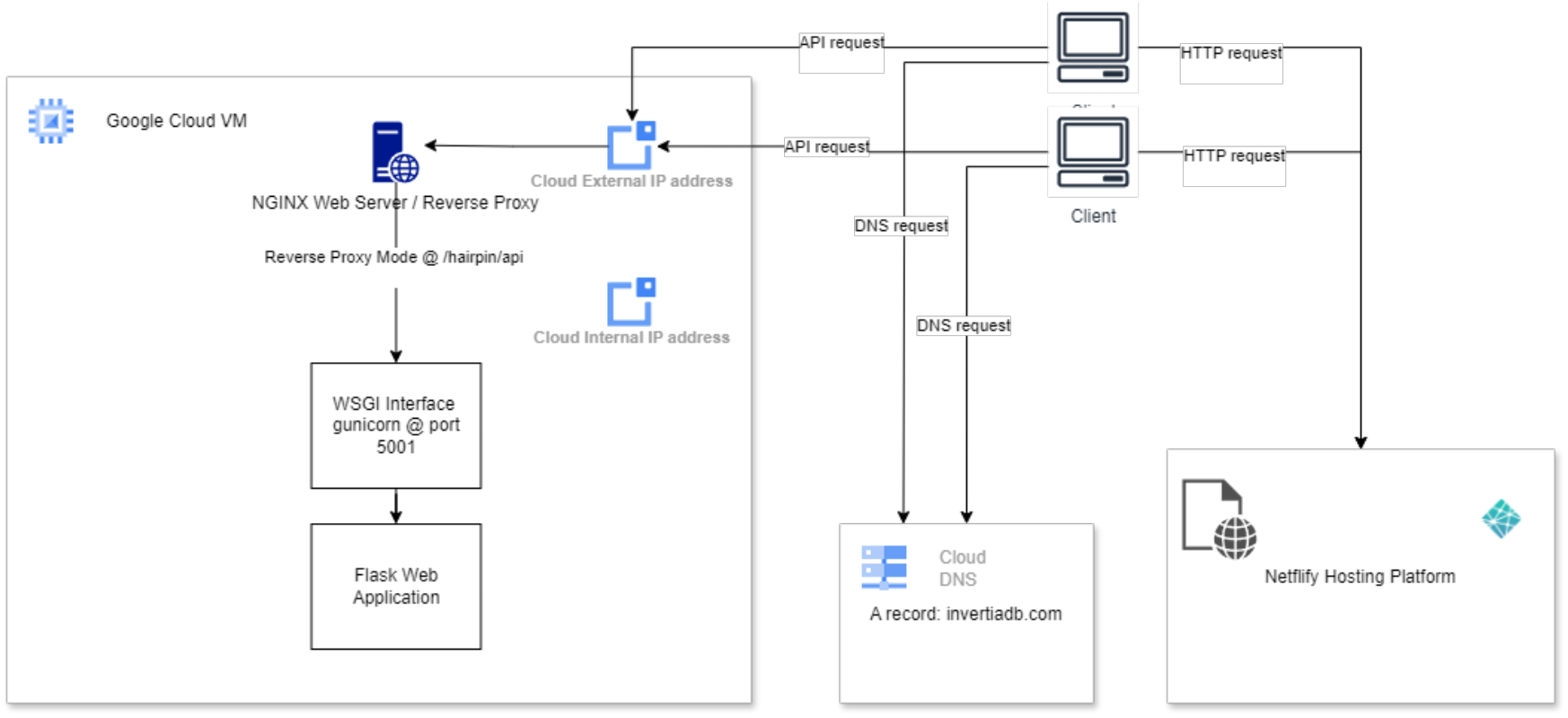
Cloud Deployment Diagram. This diagram represents the deployment of invertiaDB on the Google cloud platform.

## Notes

### Competing Interest Statement

The authors have declared no competing interest.

